# Metabarcoding malaise trap plant components enables monitoring the diversity of plant-insect interactions

**DOI:** 10.1101/2021.12.21.473674

**Authors:** Stephanie J. Swenson, Lisa Eichler, Thomas Hörren, Gerlind U. C. Lehmann, Martin Sorg, Birgit Gemeinholzer

## Abstract

The declines observed in insect abundance and diversity in the past decades has also been observed in plants, and these events are most certainly correlated. Rapid largescale biomonitoring of both plants and insects can help monitor these changes and inform decisions for land management and species protection. Malaise traps have been used for nearly 80 years for passive insect sampling of primarily flying insects, and when they enter these traps, they carry the fragments of the plants they have visited, either as plant fragments and pollen on the body surface, or as digested food material in gut contents. DNA metabarcoding is a potential method to identify these plant traces in the ethanol of the malaise bottles, which is not possible with traditional microscopy. Metabarcoding could offer more insight into what plants insects are directly interacting with at a given time, and allow for the detection of rare plants, and neophyte species visited by insects. This study, to our knowledge, is the first examination of DNA metabarcoding plant traces from Malaise trap samples, we examine 105 samples from 21 sites throughout Germany collected in a 2-week period in May of 2020. Here we report on the feasibility of sequencing these sample types, analysis of the resulting taxa, the usage of cultivated plants by insects near nature conservancy areas, and the detection of rare and neophyte species.

## Introduction

Landscape level change and chemical input in agriculture are major contributors to the rapid level of decline in diversity and abundance of insect biofauna observed in recent decades (Uhler et al. 2021). These declines are echoed in plant diversity over the past ~60 years (Eichenberg et al. 2020) and these events are most certainly linked. These rapid changes pose several risks to environmental and human health (Samways et al. 2020, van der Sluijs 2020) and necessitate improved, rapid methods of biomonitoring in order to address areas most affected by plant and insect declines, which could aid in development of best practices to alleviate these declines in natural ecosystems, while protecting economic and agricultural concerns.

Due to these declines and the fast pace of landscape changes, development of novel methods of monitoring flora in order to understand which plant resources are directly used by insects on a temporal scale, could lead to better pest management practices in agriculture as well as land management decision for scale and spacing of conservation areas. Malaise traps have been used for collection of flying insects for nearly 80years (Malaise 1937) and with higher intensity after Townes (1972) published his trap model and provide a good assessment of the flying insects in an area at a given time (Skvarla et al. 2021). When insects enter the bulk malaise trap preservative, they carry on their bodies evidence of the environments they have been living in and in their digestive tract the organisms they have been directly feeding on. The ability to identify these traces would enable a snapshot in time of the insect taxa in the area as well as the plant taxa they are interacting with or feeding on in this time period. This could enhance knowledge from traditional vegetation surveys, when they are available for an area, by providing information of which plant taxa in the environment are used on a temporal scale. Strategically placed Malaise traps could provide information on insect foraging travel to urban areas and agricultural land from more natural environments, with the detection of garden or crop species in traps placed on protected land. Additionally, identification of these traces could aid in detection of threatened plant species and encroachment of neophyte or invasive species into new areas.

In spite of this potential, identification of the pollen contained in malaise traps for would be extremely time consuming and require extensive training, and identification of plant fragments, or regurgitated food material would next to impossible with traditional microscopy, however advancing techniques in genetic identification of complex mixed species environmental samples provides a potential resource to identify these Malaise sample trap plant components. Metabarcoding, using a short gene region, or barcode, for the identification of many taxa contained in a complex sample, has displayed a great potential for plant and pollen identification over the last decade (Bell et al. 2016, Deiner et al. 2017 Hawkins et al. 2015, Ruppert et al. 2019). The application of metabarcoding to malaise trap sediments samples offers an exciting prospect for passive environmental monitoring of insects and plant interactions.

The method is not without its limitation however, and great care must be taken in barcode and primer choice, utilization of sterile techniques, extraction and PCR optimization, inclusion of several positive and negative control samples, using tested data analysis pipelines, and evaluation of the completeness of DNA reference databases used for identification (Deiner et al. 2017, Kolter and Gemeinholzer 2021). Metabarcoding, to our knowledge, has not yet been attempted with Malaise trap plant sediment and careful evaluation of the results must be performed to confirm it’s efficacy. A factor that makes this sample type different from those of previous plant or pollen metabarcoding studies is that the sample contains two signal types 1) pollen and plant material carried in on the insect body and 2) digested plant and pollen emptied from the digestive tract after entrance in to the trap. The digestion process is likely to degrade the plant DNA from the gut contents which may require amplification of shorter DNA barcoding regions. Additional consideration is required to evaluate whether these signals compete with each other, further complicating the well documented PCR bias already displayed in metabarcoding (Krehenwinkel et al. 2017). If competition is displayed barcode choice will be critical to the intention of the study (i.e., pollination studies vs. crop pest monitoring).

Passive contamination from anemophilic pollen species in flight in the duration of the collection period is an additional complicating factor, and is an unavoidable potential source of contamination for this sample type. It is likely that the occurrence of these pollens entering the collection bottle itself during the collection period will be low, however they will most certainly adhere to the outside of the insects and bottles and might enter the sample during processing steps.

This study, as part of the project Diversity of Insects in Nature Protected Areas (DINA) (Lehmann et al. 2021) aims to address whether Illumina MiSeq metabarcoding using the ITS2 plant barcode of plants fragments from Malaise trap samples can retrieve a realistic assemblage of the vegetation available and utilized by the insects found in the traps. The ITS2 barcode was chosen for its high rate of success for species level identification as well as it having one of the most abundant reference sequences available in public DNA sequencing libraries (Kolter and Gemeinholzer 2020). The source of our samples were Malaise traps placed at 21 sites throughout Germany on a gradient from internal to agriculture to internal to a nature conservancy area (Lehmann et al. 2021) for a two-week interval in May 2020. To evaluate our results, we developed the following hypotheses:

1. The majority of our samples will provide adequate data if a sufficient amount of plant data is contained in the sample.
2. Taxa retrieved will represent a realistic assemblage of German taxa, and primarily will be species with pollen available in this time period.
3. Insect will travel in to and out of conservancy areas to find food sources, and evidence of garden plant and crop plant species will be found in the internal most trap in the nature conservancy area.
4. Threatened and endangered plant species will be detected in the samples, but likely in very low quantity of reads.
5. High levels anemophilic pollen species will be observed in the samples for those species in moderate to peak pollen flight during the collection period.

## Materials and Methods

### Collection methods/sites

The collaborating project partners in the Entomological Society Krefeld (EVK) in cooperation with volunteers from the Naturschutzbund Deutschland e. V. (NABU) as well as local volunteers involved in conservation of the area established and maintained 21 sites throughout Germany (Figure 1) with five Malaise traps in each site placed at a gradient with the first located 25 M into to arable land and subsequent traps located 25 M distance from the other, the second trap located directly on the intersection of arable and protected land and the fifth located about100 M into the nature protected area (Lehmann et al 2021). Collection occurred for a duration of two weeks in the middle to end of May 2020, although exact sampling dates varied by a few days between sites. Following the collection duration, the original collection ethanol with the plant fragments was removed and saved for plant metabacording. The insects from the samples were divided into two representative samples, one for morphological identification and voucher specimens, and the other for insect metabarcoding, and DNA quality ethanol was replaced in each insect subsample.

**Figure 1:**
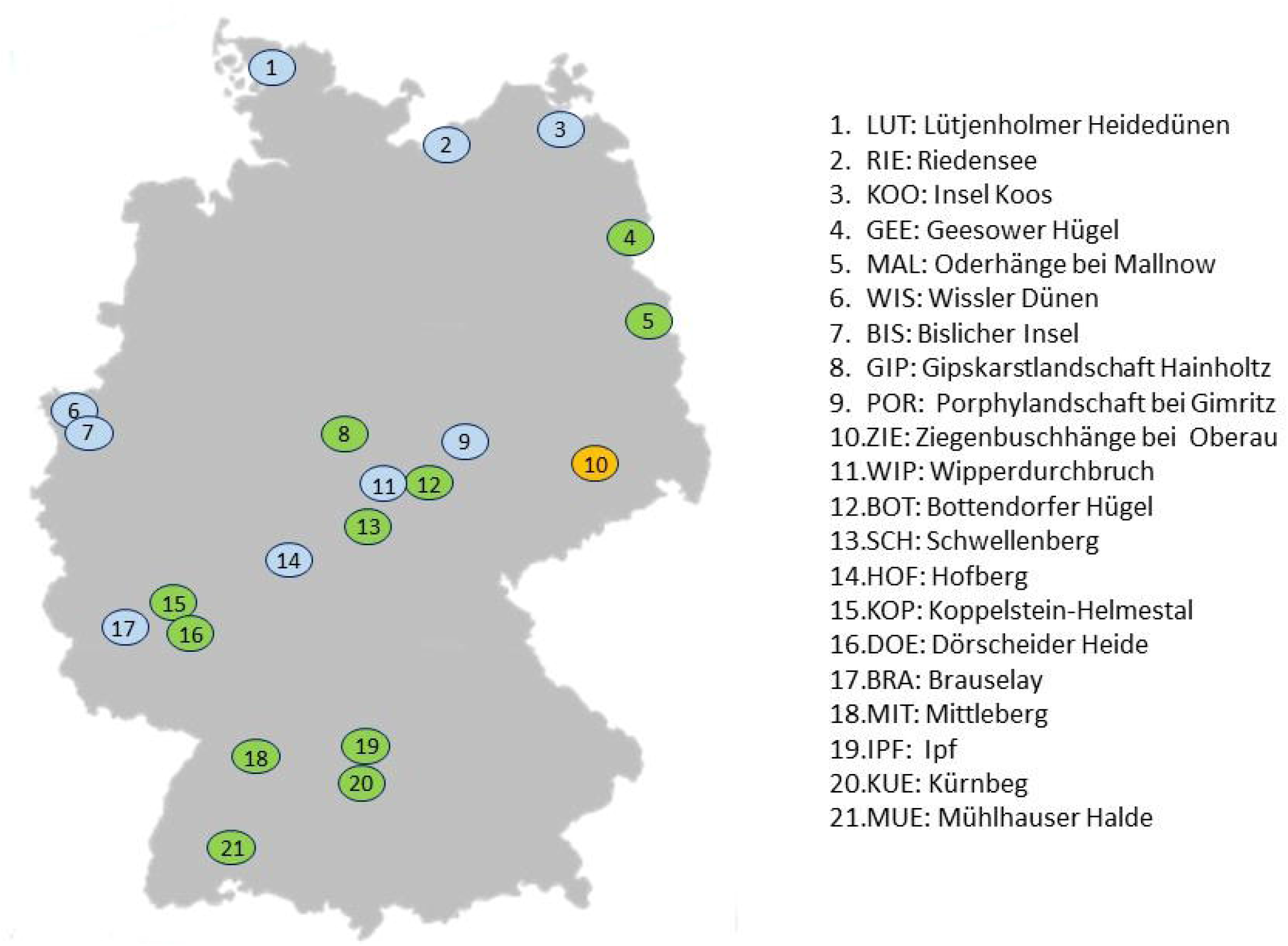
Map of the Malaise trap sampling sites throughout Germany, with site codes and names. Green indicates sites with complete data for all 5 traps, blue sites are missing data from at least 1 trap, and orange indicates no data for the site.

### Plant metabarcoding lab protocol

We vacuum filtered 200-400 mL collection ethanol using 250mL Nalgene® single use analytical filter funnels with a diameter of 47mm and 0.2 μl pore size cellulose nitrate (CN) filter in a biosafety cabinet with sterile, DNA free equipment. Following filtration, we cut the CN filter in half with a scalpel, with one half used for DNA extraction and the other saved as a voucher or backup for protocol optimization. We placed each half in a 2 mL SafeSeal microcentrifuge tube (Sarstedt AG & Co. KG) and stored them at −20°C until further processing.

We extracted DNA with Macherey Nagel Nucleomag Plant kit with the following changes: 1) to 2 mL microcentrifuge tube containing the half filter paper with sediment 1 gm of 1.4 mm ceramic beads, 500 mL MC1, 5 μl ProK, 5 μl RnaseA were added and then exposed to a 2.5 minutes duration a on Retsch MM400 bead mill at 30 hz. 2) samples were incubated with shaking at 65°C for one hour, we mixed the samples by inverting the tubes every 10 minutes 3) following incubation, we centrifuged samples for 10 minutes 4) 250-300 mL of lysate was transferred to clean tubes 5) 300 mL of MC2 and 15 mL of magnetic beads were added 6) remaining reagents were used at 25% of the standard protocol with the exception of MC6 7) 35 mL of MC6 was added and then incubated at 50°C to evaporate any residual ethanol 8) we removed 25 mL for PCR and sequencing and 2mL for DNA quantification with Qubit™ 4 fluorometer.

We performed PCR with three replicates per sample, with addition of three DNA extraction blanks, three PCR blanks, three mock community positive controls comprised of four species (*Ambrosia artemsisiifolia, Fagus sylvatica, Lilium longiflorum, Plantago lanceolata*) to validate the efficacy of laboratory protocols. We used an adaptation of the Canadian Centre for Barcoding Platinum® Taq Protocol (Ivanova and Grainger 2007) for PCR with the addition of 0.25 μL of BSA and 1.25 μL of 50% DMSO in a total volume of 12.5 μL per reaction and the ITS2 primers, Forward: ITS-3p62plF1, ACBTRGTGTGAATTGCAGRATC and Reverse: ITS-4unR1, TCCTCCGCTTATTKATATGC (Kolter and Gemeinholzer 2021). PCR cycling conditions were 95°C for 3 minutes, followed by 35 cycles of 95°C for 30 seconds, 50°C for 30 seconds, 72°C for 45 seconds, and a final extension of 72°C for 10 minutes. Following PCR cycling the three replicates were combined with 5 μL of each replicate for a total volume of 15 μL and purified with Thermo Scientific™ Exonuclease 1. The pooled replicates of non-indexed PCR products were sent to LGC Genomics GmbH where indices were added and sequencing was performed with the Illumina protocol in the 2 x 300 bp format.

### Plant metabarcoding data pipeline

We processed the sequencing data with USEARCH (Edgar 2010) and DADA2 (Callahan et al. 2016) with R (R Core Team 2020). We trimmed sequencing primers and quality filtered with a maximum expected error of 1.0 in USEARCH. We then used DADA2 for error learning, denoising by the error profile (pseudo pooling) and merging of reads. We removed Chimeras with Uchime3, and SINTAX (Edgar 2016) was used for identification with a database created in August of 2021 (A. Kolter personal communication). Obvious misidentifications were manually corrected. Average values of Amplicon Sequence Variants (ASVs) found in extraction and filter blanks were used to establish a detection threshold for significance of these ASVs occurring in samples.

## Results

### Sample evaluation

Extraction and PCR blanks were primarily fungal contaminants and contained very few reads of taxa found in the Malaise trap samples. The four species mock community positive control retrieved all species, these species were absent in the Malaise trap samples. Of the 21 sites, 11 sites produced full sequencing data for all five Malaise trap samples, 10 sites had one or more traps fail to amplify or sequencing results were primarily fungal contaminants, and one site had all traps fail to amplify (Figure 1). Overall, 80 of the 105 samples provided sequencing data.

### Plant Identification

We identified 60 families in our reads, with 218 genus level identifications, and 245 species level identifications (Supplementary table 1). The range of different families in a site was 17 (BIS) to 34 (MUE and HOF). The dominant families by percentage of total reads were Brassicaceae (34.5%, with *Brassica* sp. at 29.3%), Fabaceae (9%), Rosaceae (9%), Poaceae (8.8%), Ranunculaceae (7.8%) and Pinaceae (6.5%) and one species in Adoxaceae, *Sambucus nigra* (4%).

Species level identification *Brassica* sp. ASVs to species level was not possible with certainty, but we believe at least, *B. napus, B. rapa, B. oleracea,* and *B. nigra* to be present, and due to the prevalence in agriculture throughout Germany we believe *B. napus* to the most abundant species in the areas surrounding of experimental sites.

The highest generic level diversity was displayed in Asteraceae (20), Brassicaceae (29), Caryphyllaceae (10), Fabaceae (16), Poaceae (27), and Rosaceae (12). Of the taxa that could be assigned to at least generic level, only 6 were certain to not have pollen available in the sampling time (*Hedera* sp., *Helianthus annus, Alnus* sp., *Carpinus betulus, Corylus* sp. and *Taxus* sp.)

### Agricultural and Garden plants in Malaise traps most internal to Nature protected areas

We retrieved data for 15 Malaise traps placed most internal to the nature conservation area of the 21 sites. In these traps we detected 21 agricultural or garden plants (Table 1). Four agricultural plants were detected and were the most prevalent across the internal traps of the 15 sites. *Brassica* sp. was detected in 12 sites. Poaceae agricultural species were detected in a majority of sites (*Poa trivialis* 11 sites, *Secale cereale* 9 sites, *Triticum* sp. 7 sites) but often at low read counts that could represent false positives. 17 species of garden plants were identified in internal traps, but most often represented in low read counts which could represent false positives.

**Table 1:** Garden and Agricultural plants found in the Malaise trap samples of traps located innermost in Nature conservancy areas. Numbers indicate the number of sequencing reads retrieved.

| Taxonomy |  |  | Sites |  |  |  |  |  |  |  |  |  |  |  |  |  |  |
| --- | --- | --- | --- | --- | --- | --- | --- | --- | --- | --- | --- | --- | --- | --- | --- | --- | --- |
| Family | Genus | Species | BOT | BRA | DOE | GEE | GIP | HOF | IPF | KOO | KOP | KUE | MAL | MIT | MUE | RIE | SCH |
| **Garden plants** |  |  |  |  |  |  |  |  |  |  |  |  |  |  |  |  |  |
| Adoxaceae | *Viburnum* | *opulus* | 0 | 0 | 0 | 0 | 0 | 0 | 5 | 0 | 0 | 4 | 0 | 0 | 0 | 0 | 0 |
| Amaryllidaceae | *Allium* | *ursinum* | 0 | 0 | 0 | 0 | 164 | 0 | 0 | 0 | 0 | 0 | 0 | 0 | 0 | 0 | 0 |
| Apiaceae | *Chaerophyllum* | *roseum* | 0 | 0 | 0 | 0 | 0 | 0 | 3 | 0 | 0 | 0 | 0 | 0 | 0 | 0 | 0 |
| Apiaceae | *Heracleum* | *dissectum* | 0 | 0 | 0 | 0 | 0 | 0 | 0 | 0 | 0 | 0 | 0 | 12 | 1 | 0 | 0 |
| Asteraceae | *Achillea* | *biebersteinii* | 0 | 0 | 37 | 0 | 0 | 0 | 0 | 0 | 0 | 0 | 0 | 0 | 0 | 0 | 0 |
| Asteraceae | *Helianthus* | *annuus* | 0 | 0 | 0 | 0 | 0 | 13 | 0 | 0 | 0 | 0 | 0 | 0 | 0 | 0 | 0 |
| Asteraceae | *Pilosella* | *castellana* | 142 | 0 | 0 | 0 | 0 | 0 | 0 | 0 | 0 | 703 | 0 | 0 | 120 | 0 | 0 |
| Asteraceae | *Symphyotrichum* | *cordifolium* | 0 | 25 | 0 | 0 | 0 | 0 | 0 | 0 | 0 | 0 | 0 | 0 | 0 | 0 | 0 |
| Brassicaceae | *Aubrieta* | *olympica* | 0 | 0 | 0 | 0 | 0 | 0 | 0 | 0 | 0 | 0 | 0 | 0 | 69 | 0 | 0 |
| Brassicaceae | *Aubrieta* | sp. | 3 | 0 | 0 | 0 | 0 | 0 | 0 | 0 | 0 | 0 | 0 | 0 | 282 | 0 | 0 |
| Brassicaceae | *Aurinia* | *saxatilis* | 0 | 0 | 0 | 0 | 0 | 0 | 0 | 0 | 0 | 0 | 0 | 0 | 0 | 0 | 3 |
| Caryophyllaceae | *Cerastium* | *alpinum* | 0 | 0 | 0 | 0 | 0 | 1 | 0 | 0 | 0 | 16 | 2 | 0 | 0 | 0 | 1 |
| Cyperaceae | *Cyperus* | *diandrus* | 0 | 0 | 14 | 0 | 0 | 0 | 0 | 0 | 0 | 0 | 0 | 0 | 10 | 0 | 0 |
| Ericaceae | *Erica* | *arborea* | 0 | 0 | 0 | 0 | 0 | 0 | 0 | 0 | 0 | 0 | 0 | 34 | 0 | 0 | 0 |
| Fabaceae | *Wisteria* | sp. | 0 | 0 | 0 | 0 | 0 | 0 | 14 | 0 | 0 | 0 | 0 | 0 | 0 | 0 | 0 |
| Oleaceae | *Syringa* | *vulgaris* | 0 | 0 | 0 | 2 | 0 | 0 | 0 | 0 | 0 | 0 | 0 | 0 | 3 | 0 | 0 |
| Solanaceae | *Solanum* | *lycopersicum* | 0 | 8 | 0 | 0 | 0 | 0 | 0 | 0 | 0 | 1 | 0 | 0 | 2 | 0 | 0 |
| **Agricultural plants** |  |  |  |  |  |  |  |  |  |  |  |  |  |  |  |  |  |
| Brassicaceae | *Brassica* | sp. | 25834 | 63 | 4238 | 4524 | 3448 | 649 | 71 | 20249 | 0 | 3553 | 808 | 0 | 20 | 50605 | 1267 |
| Poaceae | *Poa* | *trivialis* | 0 | 14 | 7 | 0 | 2 | 7 | 19 | 0 | 33 | 28 | 10 | 1224 | 31 | 3 | 0 |
| Poaceae | *Secale* | *cereale* | 11 | 0 | 23 | 6 | 0 | 10 | 32 | 0 | 1 | 0 | 843 | 0 | 0 | 2 | 5 |
| Poaceae | *Triticum* | sp. | 1 | 0 | 46 | 0 | 0 | 14 | 3 | 0 | 0 | 1 | 0 | 0 | 7 | 0 | 4 |

### Plant diversity detected in the traps at all sites combined

#### Red list and neophyte taxa detection

We detected 22 Red list species (Metzing et al. 2018) in our samples (Table 2), with three Red list 2 (highly threatened), 6 Red list 3 (threatened), and 13 Red list V (nearly threated), in many cases read counts of these species were very low (under 10) and could represent false positives. 13 Neophyte species were detected in the sample, as with the Red list species, in many cases the low read counts at certain sites could indicate false positives.

**Table 2.**
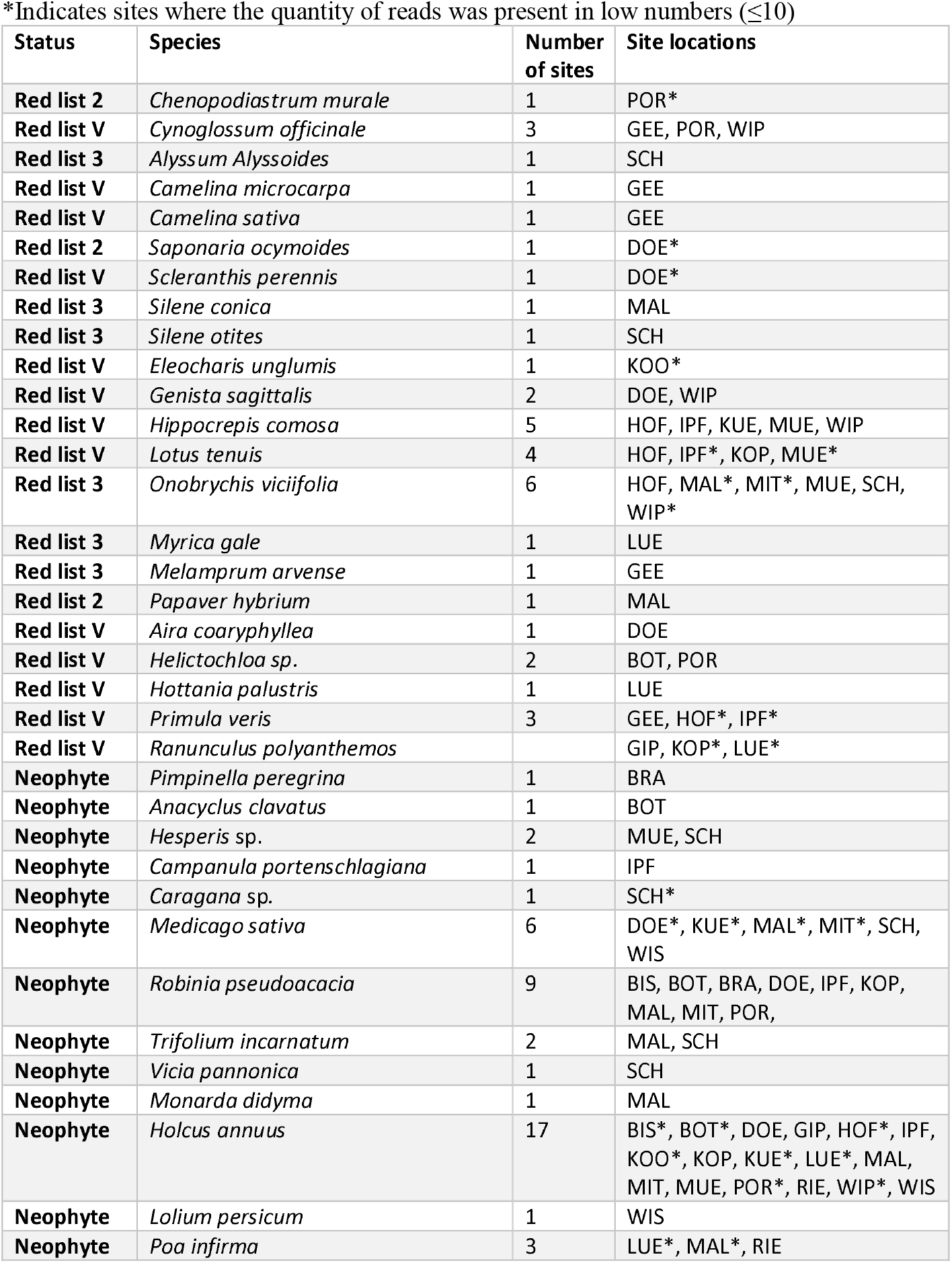
Red list plants and neophyte plant species found in the identified from ITS2 Metabarcoding reads of Malaise trap plant fragments from 21 sites throughout Germany.

#### Anemophilic pollen detection

The Stiftung Deutscher Polleninformationsdienst (PID) recorded pollens with moderate to peak flight times over the collection period as Pinaceae, Poaceae, *Rumex* sp., *Plantago* sp. *Sambucus* sp., *Quercus* sp., and *Aesculus* sp., however *Sambucus* sp., and *Aesculus* sp. are zoophilic species but they are detected in airborne pollen traps and recorded. The percentage of these taxa also occurred in relatively high quantities in our samples with the exception of *Plantago* sp. which was rarely detected: Pinaceae (6.5%), Poaceae (8.8%), *Rumex* sp. (2.3%). *Sambucus* sp. (4%), *Quercus* sp. (0.6%), and *Aesculus* sp. (0.7%)

## Discussion

### Evaluation of methods

The low number of reads of species in extraction and PCR negative controls that also appear in the Malaise trap samples indicate a very low level of contamination between samples and soundness of our laboratory protocols. Additionally, the presence of all four species contained in the positive control samples while not appearing in the Malaise trap samples, adds evidence of sound methods of extraction, primer choice, and PCR protocols. Never-the-less 25 of our 105 samples failed to amplify or were primarily non-target fungal species. Attempts to reamplify these samples with different PCR conditions have thus far failed. There are several reasons for these failures including a need for optimization of DNA extraction and/or PCR, PCR inhibiting content in the samples, plant content in the sample too low for downstream processing, mechanical disruption from high content of non-plant material in the sample (i.e., Lepidoptera wing scales), variability in ethanol concentration, and time in storage until filtration and DNA extraction (Baksay et al. 2020, Kolter and Gemeinholzer 2021, Hallmaier et al. 2018). Our results indicate that there might not be a “one size fits all” method for samples of this type that vary in collection time or duration, and environment and landscape type, and trade-off between 100% sample retrieval and high-throughput of a large number of samples may be necessary when designing a study including metabarcoding of Malaise trap plant components.

### Taxonomic retrieval

Our samples resulted in 218 genus level identifications, and 245 species level identifications from 58 families, from a 2-week collection duration in late May of 2020 in Germany. Diversity of species was highest in families with highly diverse families Asteraceae, Brassicaceae, Caryphyllaceae, Fabaceae, Poaceae, and Rosaceae, and lowest in families with low diversity in Germany.

The majority of taxa retrieved were in blossom over the collection period, and likely are the result of body pollen carried into the Malaise traps, this is in line with the malaise trap design, which is a preferred method for collection of flying insects such as the pollinator rich orders, Diptera and Hymenoptera (Brown 2021, Prado et al. 2017). We have however, with preliminary experimentation, seen evidence that the signals from fresh pollen may compete with the signals from digested plant material with the ITS2 barcode. Further investigation with mock communities and alternate barcodes is necessary to examine this possibly biased outcome.

### Occurrence of species of interest

#### Garden and Agricultural plants

We detected several species known garden plants and agricultural plant in Malaise traps located most internally in the nature protected areas, indicating travel by a proportion of insects represented in the sample into urban and agricultural landscapes for foraging. The garden species represented in these samples are mostly low abundance readings that are not relevant to insect food availability in conservation areas at this time of year, but could be indicative of insect flight distances at times of lower pollen availability. Four species of agriculture plants were found in the innermost Malaise traps. The three Poaceae species, *Poa trivialis, Secale cereale, and Triticum* sp. were present in the majority of these traps, it seems unlikely these are primarily the result of insect foraging but were inadvertently carried into the trap on the insects’ bodies. Poaceae is an anemophilic family and these plants were likely to be in pollen flight at the collection time and presence in the samples likely represents passive contamination. In addition, as with the garden plants, several occurrences occur in low reads and likely indicate false positives.

The high level of *Brassica* sp. in the innermost traps, as well as the samples as a whole, confound data interpretation. One problem being the complication in species level identifications due to polyploidy, as B. napus originated from a hybrid of *B. oleracea* and *B. rapa.* Due to its prevalence in cultivation throughout Germany and its mass flowering event occurring over this collection duration we believe the majority of reads to be *B. napus,* however we were unable to distinguish this from three other species *B. rapa, B. oleracea,* and *B. nigra* with certainty and incidence of any one of these species is indeterminate. The pollen of *B. napus* is protein rich (Borutinskaite et al., 2017) and flowering occurs as a mass event, making this an attractive food source for pollinators in the area where it is planted. Most instances of this species likely represent foraging events, however *B. napus* is also partially anemophilic so some presence is certainly due to passive contamination.

#### Red list species and Neophyte detection

We detected several threated and near threatened Red list (Metzing 2018) species in our samples (Table 2). This is an exciting prospect for biomonitoring rare plants occurrence, however due to the rareness in the environment, they are also rare in the samples and actual presence or absence needs to be carefully considered. Any incidence of threatened species in metabarcoding should be confirmed with vegetation surveys of the area before land management or protection decisions be made. We also found several neophyte species in the samples, another exciting prospect for biomonitoring, however with a different goal than protection of threatened species. Long term biomonitoring using these sample types could aid in tracking and preventing the spread of deleterious plant species.

### Passive contamination from anemophilic species

We uncovered high proportions of Anemophilic taxa in our samples, particularly Pinaceae and Poaceae, with peak pollen flight coinciding with the sampling period. This poses a problem for studies that wish to examine purely plant-insect interactions. During peak pollen it is nearly impossible to avoid all contamination from mass pollen wind pollen species, and the contamination can occur during collection, while changing and processing collection bottles, from laboratory surfaces and during filtration of the preservative ethanol. Creation of parallel blank samples in the field is likely not feasible for in terms of time and cost, however, incorporation of blanks in each of the processing phases could aid creation of thresholds and filtering species known to be at peak pollen flight.

## Conclusions

- Our study has illustrated the potential of Malaise trap plant metabarcoding as an additional tool for large scale plant biomonitoring, however cautious consideration of its limitations must be included in project design and data analysis and interpretation.
- Further experimentation needs to be undertaken to account for the reasons of sample failure with mock communities and examination of several barcodes and primer combinations.
- Passive contamination from airborne species greatly confounds data interpretation, creation of novel blanks that could be added to field protocols could aid in accounting for this source of contamination

## Acknowledgements

We are especially thankful to all the volunteers in the field who changed the collection bottles and maintained the malaise traps. Furthermore, we would like to thank all partners of the DINA consortium for the collaboration and scientific input and especially Wolfgang Wägele for project initiation. Thank you to Sabine Mutz, Kathrin Moses-Luehrsen and Annalena Kurzweil (all University Giessen) for laboratory and administrative support. Additional thanks to Andreas Kolter (University Kassel) for help with data analysis.

