## Supplementary Table 1 for "Metabarcoding malaise trap plant components enables monitoring the diversity of plant-insect interactions"

Supplementary table 1: Generic and species level identification of Amplicon Sequence Variance (ASVs) resulting from ITS2 metabarcoding of Malaise trap plant debris…

| Family | Genus | Species | Pollen available at sampling time | Zoophilic or Anemophilic | Plant status |
| --- | --- | --- | --- | --- | --- |
| Adoxaceae | *Sambucus* | sp. | YES | Zoophilic-but found in airborne traps | Wild/Native |
|  | *Sambucus* | *nigra* | YES | Zoophilic-but found in airborne traps | Wild/Native |
|  | *Viburnum* | sp. | YES | Zoophilic | Wild/Native |
|  | *Viburnum* | *opulus* | YES | Zoophilic | Garden /wild |
| Amaranthaceae | *Chenopodiastrum* | *hybridum* | YES | Zoophilic | Archeophyte/in flower mixes |
|  | *Chenopodiastrum* | *murale* | YES | Zoophilic | Red List 2 |
| Amaryllidaceae | *Allium* | sp. | YES | Zoophilic | Wild/Native |
|  | *Allium* | *aflatunenese* | YES | Zoophilic | Garden plant |
|  | *Allium* | *schoenoprasum* | YES | Zoophilic | Garden/wild |
|  | *Allium* | *stipitatum* | YES | Zoophilic | Garden plant |
|  | *Allium* | *ursinum* | YES | Zoophilic | Garden/wild |
| Apiaceae | *Anthriscus* | sp. | YES | Zoophilic | Wild/Native |
|  | *Anthriscus* | *caucalis* | YES | Zoophilic | Wild/Native |
|  | *Anthriscus* | *cerefolium* | YES | Zoophilic | Wild/Native |
|  | *Anthriscus* | *sylvestris* | YES | Zoophilic | Wild/Native |
|  | *Bunium* | *persicum* | YES | Zoophilic | Wild/Native |
|  | *Chaerophyllum* | *roseum* | YES | Zoophilic | Garden plant |
|  | *Chaerophyllum* | *temulum* | YES | Zoophilic | Wild/Native |
|  | *Conopodium* | *majus* | YES | Zoophilic | Wild/Native |
|  | *Dichoropetalum* | sp. | YES | Zoophilic | Wild/Native |
|  | *Heracleum* | *dissectum* | YES | Zoophilic | Garden plant |
|  | *Meum* | *athamanticum* | YES | Zoophilic | Wild/Native |
|  | *Pimpinella* | *saxifraga var. major* | YES | Zoophilic | Wild/Native |
|  | *Pimpinella* | *peregrina* | YES | Zoophilic | Neophyte |
|  | *Pimpinella* | *saxifraga* | YES | Zoophilic | Wild/Native |
| Araliaceae | *Hedera* | sp. | NO | Zoophilic | Wild/Native |
| Asparagaceae | *Asparagus* | *oligoclonos* | YES | Zoophilic | Garden plant |
| Asteraceae | *Achillea* | sp. | YES | Zoophilic | Wild/Native |
|  | *Achillea* | *biebersteinii* | YES | Zoophilic | Garden plant |
|  | *Anacyclus* | *clavatus* | YES | Zoophilic | Garden plant / Neophyte |
|  | *Anthemis* | sp. | YES | Zoophilic | Wild/Native |
|  | *Anthemis* | *arvensis* | YES | Zoophilic | Wild/Native |
|  | *Artemisia* | sp. | YES | Zoophilic | Wild/Native |
|  | *Bellis* | *perennis* | YES | Zoophilic | Wild/Native |
|  | *Centaurea* | sp. | YES | Zoophilic | Wild/Native |
|  | *Centaurea* | *cyanus* | YES | Zoophilic | Wild/Native |
|  | *Centaurea* | *montana* | YES | Zoophilic | Wild/Native |
|  | *Crepis* | *vesicaria* | YES | Zoophilic | Wild/Native |
|  | *Helianthus* | *annuus* | NO | Zoophilic | Agriculture/Garden/seed mixes |
|  | *Hieracium* | sp. | YES | Zoophilic | Wild/Native |
|  | *Jacobaea* | *vulgaris* | YES | Zoophilic | Wild/Native |
|  | *Leontopodium* | sp. | YES | Zoophilic | Garden plant |
|  | *Leucanthemum* | sp. | YES | Zoophilic | Wild/Native |
|  | *Matricaria* | sp. | YES | Zoophilic | Wild/Native |
|  | *Picris* | sp. | YES | Zoophilic | Wild/Native |
|  | *Pilosella* | sp. | YES | Zoophilic | Wild/Native |
|  | *Pilosella* | *castellana* | YES | Zoophilic | Garden plant |
|  | *Pilosella* | *hoppeana* | YES | Zoophilic | Wild/Native |
|  | *Senecio* | sp. | YES | Zoophilic | Wild/Native |
|  | *Senecio* | *nemorensis* | YES | Zoophilic | Wild/Native |
|  | *Senecio* | *vernalis* | YES | Zoophilic | Wild/Native |
|  | *Sonchus* | sp. | YES | Zoophilic | Wild/Native |
|  | *Symphyotrichum* | *cordifolium* | YES | Zoophilic | Garden plant |
|  | *Taraxacum* | sp. | YES | Zoophilic | Wild/Native |
|  | *Tragopogon* | sp. | YES | Zoophilic | Wild/Native |
|  | *Tripleurospermum* | *maritimum* | YES | Zoophilic | Wild/Native |
| Betulaceae | *Alnus* | sp. | NO | Anemophilic | Wild/Native |
|  | *Betula* | sp. | YES | Anemophilic | Wild/Native |
|  | *Carpinus* | *betulus* | NO | Anemophilic | Wild/Native |
|  | *Corylus* | sp. | NO | Anemophilic | Wild/Native |
| Boraginaceae | *Aegonychon* | *purpurocaeruleum* | YES | Zoophilic | Garden plant |
|  | *Anchusa* | sp. | YES | Zoophilic | Wild/Native |
|  | *Borago* | *officinalis* | YES | Zoophilic | Wild/Native |
|  | *Buglossoides* | *arvensis* | YES | Zoophilic | Wild/Native |
|  | *Cynoglossum* | *officinale* | YES | Zoophilic | Red list V |
|  | *Echium* | *vulgare* | YES | Zoophilic | Wild/Native |
|  | *Lithospermum* | sp. | YES | Zoophilic | Wild/Native |
|  | *Myosotis* | sp. | YES | Zoophilic | Wild/Native |
|  | *Myosotis* | *arvensis* | YES | Zoophilic | Wild/Native |
|  | *Myosotis* | *laxa* | YES | Zoophilic | Wild/Native |
|  | *Myosotis* | *sylvatica* | YES | Zoophilic | Wild/Native |
|  | *Symphytum* | *officinale* | YES | Zoophilic | Wild/Native |
| Brassicaceae | *Alliaria* | *petiolata* | YES | Zoophilic | Wild/Native |
|  | *Alyssum* | sp. | YES | Zoophilic | Wild/Native |
|  | *Alyssum* | *alysssoides* | YES | Zoophilic | Red list 3 |
|  | *Arabidopsis* | *arenosa* | YES | Zoophilic | Wild/Native |
|  | *Arabis* | sp. | YES | Zoophilic | Wild/Native |
|  | *Aubrieta* | sp. | YES | Zoophilic | Garden plant |
|  | *Aubrieta* | *olympica* | YES | Zoophilic | Garden plant |
|  | *Aurinia* | *saxatilis* | Yes | Zoophilic | Garden plant |
|  | *Barbarea* | *vulgaris* | YES | Zoophilic | Wild/Native |
|  | *Brassica* | sp. | YES | Zoophilic -but found in airborne traps | Agricultural/Wild |
|  | *Bunias* | *orientalis* | YES | Zoophilic | Wild/Native |
|  | *Camelina* | *microcarpa* | YES | Zoophilic | Red list V |
|  | *Camelina* | *sativa* | YES | Zoophilic | Red list V |
|  | *Capsella* | *bursa-pastoris* | YES | Zoophilic | Wild/Native |
|  | *Capsella* | sp. | YES | Zoophilic | Wild/Native |
|  | *Cardamine* | sp. | YES | Zoophilic | Wild/Native |
|  | *Cardamine* | *flexuosa* | YES | Zoophilic | Wild/Native |
|  | *Cardamine* | *impatiens* | YES | Zoophilic | Wild/Native |
|  | *Cardamine* | *pratensis* | YES | Zoophilic | Wild/Native |
|  | *Descurainia* | *sophia* | YES | Zoophilic | Wild/Native |
|  | *Diplotaxis* | *tenuifolia* | YES | Zoophilic | Wild/Native |
|  | *Draba* | *verna* | YES | Zoophilic | Wild/Native |
|  | *Ery~~s~~simum* | sp. | YES | Zoophilic | Wild/Native |
|  | *Erysimum* | *cheiri* | YES | Zoophilic | Wild/Native |
|  | *Hesperis* | sp. | YES | Zoophilic | Garden plant/neophyte |
|  | *Iberis* | *sempervirens* | YES | Zoophilic | Garden plant |
|  | *Isatis* | sp. | YES | Zoophilic | Wild/Native |
|  | *Lepidium* | sp. | YES | Zoophilic | Wild/Native |
|  | *Lepidium* | *campestre* | YES | Zoophilic | Wild/Native |
|  | *Lepidium* | *draba* | YES | Zoophilic | Wild/Native |
|  | *Microthlaspis* | *perfoliatum* | YES | Zoophilic | Wild/Native |
|  | *Nasturtium* | *officinale* | YES | Zoophilic | Wild/Native |
|  | *Raphanus* | *sativus* | YES | Zoophilic | Wild/Native |
|  | *Rorippa* | sp. | YES | Zoophilic | Wild/Native |
|  | *Sinapis* | *alba* | YES | Zoophilic | Wild/Native |
|  | *Sisymbrium* | *loeselii* | YES | Zoophilic | Wild/Native |
|  | *Sisymbrium* | *officinale* | YES | Zoophilic | Wild/Native |
|  | *Teesdalia* | *nudicaulis* | YES | Zoophilic | Wild/Native |
|  | *Thlaspi* | *arvense* | YES | Zoophilic | Wild/Native |
|  | *Turritis* | *glabra* | YES | Zoophilic | Wild/Native |
| Campanulaceae | *Campanula* | *persicifolia* | YES | Zoophilic | Wild/Native |
|  | *Campanula* | *portenschlagiana* | YES | Zoophilic | Garden plant/Neophyte |
|  | *Campanula* | *rapunculus* | YES | Zoophilic | Wild/Native |
| Caprifoliaceae | *Knautia* | sp. | YES | Zoophilic | Wild/Native |
| Caryophllaceae | *Arenaria* | *serpyllifolia* | YES | Zoophilic | Wild/Native |
|  | *Cerastium* | sp. | YES | Zoophilic | Wild/Native |
|  | *Cerastium* | *alpinum* | YES | Zoophilic | Garden plant |
|  | *Dianthus* | sp. | YES | Zoophilic | Wild/Native |
|  | *Moehringia* | *trinervia* | YES | Zoophilic | Wild/Native |
|  | *Saponaria* | *ocymoides* | YES | Zoophilic | Red list 2 |
|  | *Scleranthus* | *perennis* | YES | Zoophilic | Red List V |
|  | *Silene* | sp. | YES | Zoophilic | Wild/Native |
|  | *Silene* | *conica* | YES | Zoophilic | Red List 3 |
|  | *Silene* | *flos-cuculi* | YES | Zoophilic | Wild/Native |
|  | *Silene* | *latifolia* | YES | Zoophilic | Wild/Native |
|  | *Silene* | *otites* | YES | Zoophilic | Red List 3 |
|  | *Spergularia* | sp. | YES | Zoophilic | Wild/Native |
|  | *Stellaria* | *holostea* | YES | Zoophilic | Wild/Native |
|  | *Stellaria* | *media* | YES | Zoophilic | Wild/Native |
|  | *Viscaria* | *atropurpurea* | YES | Zoophilic | Wild/Native |
| Celastraceae | *Euonymus* | sp. | YES | Zoophilic | Wild/Native |
|  | *Euonymus* | *europaeus* | YES | Zoophilic | Wild/Native |
| Cistaceae | *Helianthemum* | sp. | YES | Zoophilic | Wild/Native |
| Convolvulaceae | *Convolvulus* | *arvensis* | YES | Zoophilic | Wild/Native |
| Cornaceae | *Cornus* | sp. | YES | Zoophilic | Wild/Native |
|  | *Cornus* | *sanguinea* | YES | Zoophilic | Wild/Native |
| Crassulaceae | *Phedimus* | *hybridus* | YES | Zoophilic | Garden plant |
| Cucurbitaceae | *Bryonia* | sp. | YES | Zoophilic | Wild/Native |
| Cupressaceae | *Juniperus* | sp. | YES | Anemophilic | Wild/Native |
| Cyperaceae | *Carex* | sp. | YES | Zoophilic | Wild/Native |
|  | *Carex* | *areniaria* | YES | Zoophilic | Wild/Native |
|  | *Carex* | *flacca* | YES | Zoophilic | Wild/Native |
|  | *Carex* | *hirta* | YES | Zoophilic | Wild/Native |
|  | *Carex* | *spicata* | YES | Zoophilic | Wild/Native |
|  | *Cyperus* | *diandrus* | YES | Zoophilic | Garden plant |
|  | *Eleocharis* | *uniglumis* | YES | Zoophilic | Red List V |
| Ericaceae | *Erica* | *arborea* | YES | Zoophilic | Garden plant |
|  | *Rhododendron* | sp. | YES | Zoophilic | Garden plant |
| Euphorbiaceae | *Euphorbia* | sp. | YES | Zoophilic | Wild/Native |
|  | *Euphorbia* | *amygdaloides* | YES | Zoophilic | Wild/Native |
| Fabaceae | *Anthyllis* | *vulneraria* | YES | Zoophilic | Wild/Native |
|  | *Caragana* | sp. | YES | Zoophilic | Neophyte |
|  | *Cytisus* | sp. | YES | Zoophilic | Wild/Native |
|  | *Securigera* | sp. | YES | Zoophilic | Wild/Native |
|  | *Genista* | *sagittalis* | YES | Zoophilic | Red List V |
|  | *Hippocrepis* | sp. | YES | Zoophilic | Wild/Native |
|  | *Hippocrepis* | *comosa* | YES | Zoophilic | Red List V |
|  | *Lathyrus* | *pratensis* | YES | Zoophilic | Wild/Native |
|  | *Lotus* | sp. | YES | Zoophilic | Wild/Native |
|  | *Lotus* | *corniculatus* | YES | Zoophilic | Wild/Native |
|  | *Lotus* | *tenuis* | YES | Zoophilic | Red list V |
|  | *Lotus* | *uliginosus* | YES | Zoophilic | Wild/Native |
|  | *Medicago* | *lupulina* | YES | Zoophilic | Wild/Native |
|  | *Medicago* | *sativa* | YES | Zoophilic | Neophyte |
|  | *Onobrychis* | sp. | YES | Zoophilic | Wild/Native |
|  | *Onobrychis* | *viciifolia* | YES | Zoophilic | Red List 3 |
|  | *Ononis* | *repens* | YES | Zoophilic | Wild/Native |
|  | *Ononis* | *spinosa* | YES | Zoophilic | Wild/Native |
|  | *Oxytropis* | sp. | YES | Zoophilic | Wild/Native |
|  | *Pisum* | *sativum* | YES | Zoophilic | Garden plant |
|  | *Robinia* | *pseudoacacia* | YES | Zoophilic | Neophyte |
|  | *Trifolium* | *arvense* | YES | Zoophilic | Wild/Native |
|  | *Trifolium* | *dubium* | YES | Zoophilic | Wild/Native |
|  | *Trifolium* | *incarnatum* | YES | Zoophilic | Neophyte/Seed mixes |
|  | *Trifolium* | *pratense* | YES | Zoophilic | Wild/Native |
|  | *Trifolium* | *repens* | YES | Zoophilic | Wild/Native |
|  | *Vicia* | sp. | YES | Zoophilic | Wild/Native |
|  | *Vicia* | *faba* | YES | Zoophilic | Agricultural/Garden plant |
|  | *Vicia* | *hirsuta* | Yes | Zoophilic | Wild/Native |
|  | *Vicia* | *pannonica* | YES | Zoophilic | Neophyte |
|  | *Vicia* | *sativa* | YES | Zoophilic | Wild/Native |
|  | *Vicia* | *sepium* | YES | Zoophilic | Wild/Native |
|  | *Vicia* | *villosa* | YES | Zoophilic | Wild/Native |
|  | *Wisteria* | sp. | YES | Zoophilic | Garden plant |
| Fagaceae | *Quercus* | sp. | YES | Anemophilic | Wild/Native |
| Geraniaceae | *Erodium* | *cicutarium* | YES | Zoophilic | Wild/Native |
|  | *Geranium* | *robertianum* | YES | Zoophilic | Wild/Native |
| Hydrangeaceae | *Deutzia* | sp. | YES | Zoophilic | Garden plant |
|  | *Deutzia* | *gracilis* | Yes | Zoophilic | Garden plant |
|  | *Deutzia* | *sieboldiana* | YES | Zoophilic | Garden plant |
|  | *Hydrangea* | *serrata* | YES | Zoophilic | Garden plant |
|  | *Philadelphus* | sp. | Yes | Zoophilic | Garden plant |
|  | *Philadelphus* | *pubescens* | YES | Zoophilic | Garden plant |
| Hydrophyllaceae | *Phacelia* | sp. | YES | Zoophilic | Flower seed mixes |
|  | *Phacelia* | *tanacetifolia* | Yes | Zoophilic | Flower seed mixes |
| Juglandaceae | *Juglans* | sp. | YES | Anemophilic | Wild/Native |
| Juncaceae | *Luzula* | sp. | YES | Anemophilic | Wild/Native |
| Lamiaceae | *Ajuga* | *reptans* | YES | Zoophilic | Wild/Native |
|  | *Glechoma* | *hederacea* | YES | Zoophilic | Wild/Native |
|  | *Lamium* | sp. | YES | Zoophilic | Wild/Native |
|  | *Lamium* | *galeobdolon* | YES | Zoophilic | Wild/Native |
|  | *Lamium* | *maculatum* | YES | Zoophilic | Wild/Native |
|  | *Monarda* | *didyma* | YES | Zoophilic | Garden plant/Neophyte |
|  | *Salvia* | sp. | YES | Zoophilic | Wild/Native |
|  | *Stachys* | sp. | Yes | Zoophilic | Wild/Native |
| Malvaceae | *Tilia* | sp. | YES | Anemophilic | Wild/Native |
|  | *Tilia* | *platyphyllos* | YES | Anemophilic | Wild/Native |
|  | *Tilia* | *tomentosa* | YES | Anemophilic | Wild/Native |
| Myricaceae | *Myrica* | *gale* | YES | Zoophilic | Red list 3 |
| Oleaceae | *Forsythia* | *suspensa* | Yes | Zoophilic | Garden plant |
|  | *Fraxinus* | *excelsior* | YES | Zoophilic | Wild/Native |
|  | *Fraxinus* | *ornus* | YES | Zoophilic | Wild/Native |
|  | *Fraxinus* | sp. | YES | Zoophilic | Wild/Native |
|  | *Ligustrum* | *vulgare* | YES | Zoophilic | Wild/Native |
|  | *Syringa* | *vulgaris* | Yes | Zoophilic | Garden plant |
| Onagraceae | *Epilobium* | *lanceolatum* | YES | Zoophilic | Wild/Native |
| Orobanchaceae | *Melampyrum* | *arvense* | Yes | Zoophilic | Red list 3 |
|  | *Rhinanthus* | *alectorolophus* | YES | Zoophilic | Wild/Native |
| Paeoniaceae | *Paeonia* | sp. | YES | Zoophilic | Garden plant |
|  | *Paeonia* | *lactiflora* | YES | Zoophilic | Garden plant |
| Papaveraceae | *Chelidonium* | *majus* | YES | Zoophilic | Wild/Native |
|  | *Papaver* | sp. | YES | Zoophilic | Wild/Native |
|  | *Papaver* | *argemone* | Yes | Zoophilic | Wild/Native |
|  | *Papaver* | *bracteatum var. psuedoorientale* | YES | Zoophilic | Garden plant |
|  | *Papaver* | *dubium* | YES | Zoophilic | Wild/Native |
|  | *Papaver* | *hybridum* | YES | Zoophilic | Red list 2 |
|  | *Papaver* | *rhoeas* | Yes | Zoophilic | Wild/Native |
| Pinaceae | *Abies* | sp. | YES | Anemophilic | Wild/Native |
|  | *Picea* | sp. | YES | Anemophilic | Wild/Native |
|  | *Pinus* | sp. | YES | Anemophilic | Wild/Native |
|  | *Pinus* | *sylvestris* | YES | Anemophilic | Wild/Native |
| Plantaginaceae | *Plantago* | sp. | YES | Anemophilic | Wild/Native |
|  | *Plantago* | *major* | YES | Anemophilic | Wild/Native |
|  | *Plantago* | *media* | YES | Anemophilic | Wild/Native |
|  | *Plantago* | *ovata* | YES | Anemophilic | Wild/Native |
|  | *Veronica* | sp. | YES | Zoophilic | Wild/Native |
|  | *Veronica* | *arvensis* | YES | Zoophilic | Wild/Native |
|  | *Veronica* | *beccabunga* | YES | Zoophilic | Wild/Native |
|  | *Veronica* | *chamaedrys* | YES | Zoophilic | Wild/Native |
|  | *Veronica* | *persica* | YES | Zoophilic | Wild/Native |
|  | *Veronica* | *polita* | YES | Zoophilic | Wild/Native |
|  | *Veronica* | *serpyllifolia* | YES | Zoophilic | Wild/Native |
| Plumbaginaceae | *Armeria* | sp. | YES | Zoophilic | Wild/Native |
| Poaceae | *Agrostis* | *capillaris* | YES | Anemophilic | Wild/Native |
|  | *Aira* | *caryphyllea* | YES | Anemophilic | Red List V |
|  | *Alopecurus* | sp. | YES | Anemophilic | Wild/Native |
|  | *Alopecurus* | *aequalis* | YES | Anemophilic | Wild/Native |
|  | *Alopecurus* | *myosuroides* | YES | Anemophilic | Wild/Native |
|  | *Alopecurus* | *pratensis* | YES | Anemophilic | Wild/Native |
|  | *Anthoxanthum* | sp. | YES | Anemophilic | Wild/Native |
|  | *Anthoxanthum* | *aristatum* | YES | Anemophilic | Wild/Native |
|  | *Anthoxanthum* | *nipponicum* | YES | Anemophilic | Wild/Native |
|  | *Arrhenatherum* | sp. | YES | Anemophilic | Wild/Native |
|  | *Arrhenatherum* | *elatius* | YES | Anemophilic | Wild/Native |
|  | *Avena* | sp. | YES | Anemophilic | Wild/Native |
|  | *Deschampsia* | *flexuosa* | YES | Anemophilic | Wild/Native |
|  | *Helictotrichon* | *pubescens* | YES | Anemophilic | Wild/Native |
|  | *Briza* | *media* | YES | Anemophilic | Wild/Native |
|  | *Bromus* | *erectus* | YES | Anemophilic | Wild/Native |
|  | *Bromus* | *hordeaceus* | YES | Anemophilic | Wild/Native |
|  | *Bromus* | *tectorum* | YES | Anemophilic | Wild/Native |
|  | *Cynosurus* | *cristatus* | YES | Anemophilic | Wild/Native |
|  | *Dactylis* | *glomerata* | YES | Anemophilic | Wild/Native |
|  | *Elymus* | *repens* | YES | Anemophilic | Wild/Native |
|  | *Festuca* | sp. | YES | Anemophilic | Wild/Native |
|  | *Festuca* | *filiformis* | YES | Anemophilic | Wild/Native |
|  | *Festuca* | *rubra* | YES | Anemophilic | Wild/Native |
|  | *Helictochloa* | sp. | YES | Anemophilic | Red list V |
|  | *Holcus* | *annuus* | YES | Anemophilic | Neophyte |
|  | *Holcus* | *lanatus* | YES | Anemophilic | Wild/Native |
|  | *Hordeum* | *murinum* | YES | Anemophilic | Wild/Native |
|  | *Hordeum* | *vulgare* | YES | Anemophilic | Agricultural plant |
|  | *Koeleria* | sp. | YES | Anemophilic | Wild/Native |
|  | *Lolium* | sp. | YES | Anemophilic | Wild/Native |
|  | *Lolium* | *canariense* | YES | Anemophilic | Garden plant |
|  | *Lolium* | *perenne* | YES | Anemophilic | Wild/Native |
|  | *Lolium* | *persicum* | YES | Anemophilic | Neophyte |
|  | *Melica* | *uniflora* | YES | Anemophilic | Wild/Native |
|  | *Milium* | *effusum* | YES | Anemophilic | Wild/Native |
|  | *Phleum* | *pratense* | YES | Anemophilic | Wild/Native |
|  | *Phleum* | sp. | YES | Anemophilic | Wild/Native |
|  | *Poa* | sp. | YES | Anemophilic | Wild/Native |
|  | *Poa* | *annua* | YES | Anemophilic | Wild/Native |
|  | *Poa* | *infirma* | YES | Anemophilic | Neophyte |
|  | *Poa* | *pratensis* | YES | Anemophilic | Wild/Native |
|  | *Poa* | *trivialis* | YES | Anemophilic | Agricultural plant |
|  | *Secale* | *cereale* | YES | Anemophilic | Agricultural plant |
|  | *Trisetum* | sp. | YES | Anemophilic | Wild/Native |
|  | *Trisetum* | *flavescens* | YES | Anemophilic | Wild/Native |
|  | *Triticum* | sp. | YES | Anemophilic | Agricultural plant |
|  | *Zea* | mays | YES | Anemophilic | Agricultural plant |
| Polygalaceae | *Polygala* | comosa | YES | Zoophilic | Wild/Native |
|  | *Polygonum* | sp. | YES | Zoophilic | Wild/Native |
|  | *Rumex* | sp. | YES | Zoophilic | Wild/Native |
|  | *Rumex* | *acetosa* | YES | Zoophilic | Wild/Native |
|  | *Rumex* | *acetosella* | YES | Zoophilic | Wild/Native |
|  | *Rumex* | *lapponicus* | YES | Zoophilic | Wild/Native |
| Primulaceae | *Hottonia* | *palustris* | YES | Zoophilic | Red list V |
|  | *Primula* | *veris* | YES | Zoophilic | Red list V |
| Ranunculaceae | *Anemone* | *nemorosa* | YES | Zoophilic | Wild/Native |
|  | *Anemone* | *sylvestris* | YES | Zoophilic | Wild/Native |
|  | *Aquilegia* | sp. | YES | Zoophilic | Garden plant |
|  | *Clematis* | sp. | YES | Zoophilic | Wild/Native |
|  | *Eranthis* | *cilicica* | YES | Zoophilic | Garden plant |
|  | *Eranthis* | *hyemalis* | YES | Zoophilic | Wild/Native |
|  | *Ficaria* | *verna* | YES | Zoophilic | Wild/Native |
|  | *Ranunculus* | sp. | YES | Zoophilic | Wild/Native |
|  | *Ranunculus* | *bulbosus* | YES | Zoophilic | Wild/Native |
|  | *Ranunculus* | *flammula* | YES | Zoophilic | Wild/Native |
|  | *Ranunculus* | *polyanthemos* | YES | Zoophilic | Red list V |
|  | *Ranunculus* | *repens* | YES | Zoophilic | Wild/Native |
|  | *Thalictrum* | sp. | YES | Zoophilic | Wild/Native |
| Resedaceae | *Reseda* | *lutea* | YES | Zoophilic | Wild/Native |
|  | *Reseda* | *luteola* | YES | Zoophilic | Wild/Native |
| Rhamnaceae | *Frangula* | *alnus* | YES | Zoophilic | Wild/Native |
|  | *Rhamnus* | sp. | YES | Zoophilic | Wild/Native |
| Rosaceae | *Alchemilla* | sp. | Yes | Zoophilic | Wild/Native |
|  | *Crataegus* | sp. | YES | Zoophilic | Wild/Native |
|  | *Filipendula* | *vulgaris* | YES | Zoophilic | Wild/Native |
|  | *Fragaria* | *viridis* | YES | Zoophilic | Wild/Native |
|  | *Geum* | sp. | YES | Zoophilic | Wild/Native |
|  | *Geum* | *rivale* | YES | Zoophilic | Wild/Native |
|  | *Kerria* | *japonica* | YES | Zoophilic | Garden plant |
|  | *Potentilla* | sp. | YES | Zoophilic | Wild/Native |
|  | *Potentilla* | *argentea* | YES | Zoophilic | Wild/Native |
|  | *Potentilla* | *erecta* | YES | Zoophilic | Wild/Native |
|  | *Potentilla* | *reptans* | YES | Zoophilic | Wild/Native |
|  | *Prunus* | sp. | YES | Zoophilic | Wild/Native |
|  | *Rosa* | sp. | YES | Zoophilic | Wild/Native |
|  | *Rubus* | sp. | YES | Zoophilic | Wild/Native |
|  | *Rubus* | *caesius* | YES | Zoophilic | Wild/Native |
|  | *Rubus* | *idaeus* | Yes | Zoophilic | Wild/Native |
|  | *Sanguisorba* | sp. | YES | Zoophilic | Wild/Native |
|  | *Sorbus* | *aucuparia* | YES | Zoophilic | Wild/Native |
| Rubiaceae | *Galium* | sp. | YES | Zoophilic | Wild/Native |
|  | *Galium* | *mollugo* | YES | Zoophilic | Wild/Native |
|  | *Galium* | *odoratum* | YES | Zoophilic | Wild/Native |
|  | *Galium* | *verum* | YES | Zoophilic | Wild/Native |
| Salicaceae | *Populus* | sp. | YES | Zoophilic | Wild/Native |
|  | *Salix* | sp. | YES | Zoophilic | Wild/Native |
| Sapindaceae | *Acer* | sp. | YES | Zoophilic | Wild/Native |
|  | *Acer* | *campestre* | YES | Zoophilic | Wild/Native |
|  | *Acer* | *platanoides* | YES | Zoophilic | Wild/Native |
|  | *Acer* | *pseudoplatanus* | YES | Zoophilic | Wild/Native |
|  | *Aesculus* | *hippocastanum* | YES | Combination | Wild/Native |
| Saxifragaceae | *Saxifraga* | sp. | YES | Zoophilic | Wild/Native |
| Scrophulariaceae | *Scrophularia* | sp. | YES | Zoophilic | Wild/Native |
| Solanaceae | *Lycium* | sp. | YES | Zoophilic | Wild/Native |
|  | *Solanum* | sp. | YES | Zoophilic | Wild/Native |
|  | *Solanum* | *dulcamara* | YES | Zoophilic | Wild/Native |
|  | *Solanum* | *lyscopersicum* | YES | Zoophilic | Garden plant |
|  | *Solanum* | *tuberosum* | Yes | Zoophilic | Agricultural/Garden plant |
| Taxaceae | *Taxus* | sp. | NO | Zoophilic | Wild/Native |
| Thesiaceae | *Thesium* | sp. | Uncertain | Zoophilic | Cannot be determined at Genus level |
| Ulmaceae | *Ulmus* | sp. | YES | Anemophilic | Wild/Native |
| Urticaceae | *Parietaria* | *judaica* | YES | Anemophilic | Wild/Native |
|  | *Urtica* | *dioica* | YES | Anemophilic | Wild/Native |
|  | *Urtica* | *urens* | YES | Anemophilic | Wild/Native |
| Violaceae | *Viola* | sp. | YES | Zoophilic | Wild/Native |
|  | *Viola* | *hirta* | YES | Zoophilic | Wild/Native |
| Vitaceae | *Vitis* | sp. | YES | Zoophilic | Agricultural plant |
